# Ultrasound imaging of in situ transcriptional activity in opaque tissue

**DOI:** 10.1101/2025.07.06.663365

**Authors:** Shirin Shivaei, Kathy Y.M. Cheung, Akanksha Yadav, Isabella U. Hurvitz, Sunho Lee, Julio Revilla, Claire Rabut, Ernesto Criado-Hidalgo, Raymond J. Zhang, Mikhail G. Shapiro

**Affiliations:** Division of Biology and Biological Engineering, California Institute of Technology; Pasadena, CA 91125, USA; Division of Chemistry and Chemical Engineering, California Institute of Technology; Pasadena, CA 91125, USA; Andrew and Peggy Cherng Department of Medical Engineering, California Institute of Technology; Pasadena, CA 91125, USA; Howard Hughes Medical Institute; Pasadena, CA 91125, USA

## Abstract

Ultrasound imaging and acoustic reporter genes provide unique capabilities for in vivo biological imaging by leveraging ultrasound’s ability to visualize opaque tissues with high spatiotemporal resolution. But until now, the expression of acoustic reporter genes – based on gas vesicle (GV) proteins – has been limited to ex vivo-modified cells due to the complexity of the GV gene cluster, precluding valuable in situ applications. Here, we develop a system capable of introducing GV genes directly into native tissues via stoichiometric multi-AAV delivery. We validate this system in the mouse brain, demonstrating well-tolerated in situ gene expression and repeated ultrasound imaging over more than a month in the same animal. Furthermore, by placing GV genes under the control of immediate early gene promoters, we demonstrate the ability to track in vivo gene expression changes arising from elevated neural activity during epileptic seizures. This work connects ultrasound to in situ transcriptional dynamics happening inside the opaque tissues of living creatures.

## INTRODUCTION

Studying biological function in intact organisms requires methods that visualize cellular activities such as gene expression in opaque tissues. In this context, established optical methods have limited penetration depth due to light scattering and absorption, while non-contact modalities such as magnetic resonance imaging or positron emission tomography lack cellular-scale resolution and require costly equipment or radioactive tracers^1^.

In contrast, ultrasound combines high spatial resolution—on the order of tens to hundreds of micrometers^2^—with centimeter-scale penetration while being relatively affordable and accessible. Recently, acoustic reporter genes (ARGs) based on gas vesicle (GV) proteins have expanded ultrasound’s potential for molecular and cellular imaging. GVs, air-filled protein nanostructures derived from photosynthetic microbes, have been established as genetically encodable labels in mammalian cells^3–5^, and engineered enzyme- and ion-sensing GV variants have begun to connect ultrasound contrast to molecular signals^6,7^. Despite these advances, one major limitation of GV-based ARGs is that they have yet to be adapted for in situ imaging of endogenous cells, such as neurons, or the transcriptional responses arising from their function. This challenge stems from the complexity of GV expression, which requires polycistronic co-expression of multiple genes at specific ratios. In fact, all previous work on ARGs has involved genetically modifying and sorting cells ex vivo, and only then injecting them into the body.

Here, we overcome this limitation by developing a system for in situ expression of GVs using stoichiometric multi-AAV delivery. We first test this system in vitro in primary cultured neurons, then validate it in vivo in the mouse brain. Following intracranial delivery, we demonstrate—for the first time— that ultrasound can image in situ gene expression in the brain of live mice in both 2D and 3D, and that this imaging can be performed repeatedly over 6 weeks. We then engineer gene circuits to place GV expression downstream of immediate-early gene (IEG) promoters, allowing ultrasound to visualize activity-induced changes in gene expression. We validate this system by imaging increased IEG transcription in the hippocampus resulting from drug-induced epileptic seizures. These in vivo results quantitatively correlate with subsequent post-mortem histology. By leveraging ultrasound’s ability to image deep tissues, this platform enables longitudinal, non-terminal imaging of transcriptional activity in the same animal (**Fig. 1**), a significant improvement over terminal histological methods and optical imaging techniques limited by invasiveness and tissue opacity.

**Figure 1:**
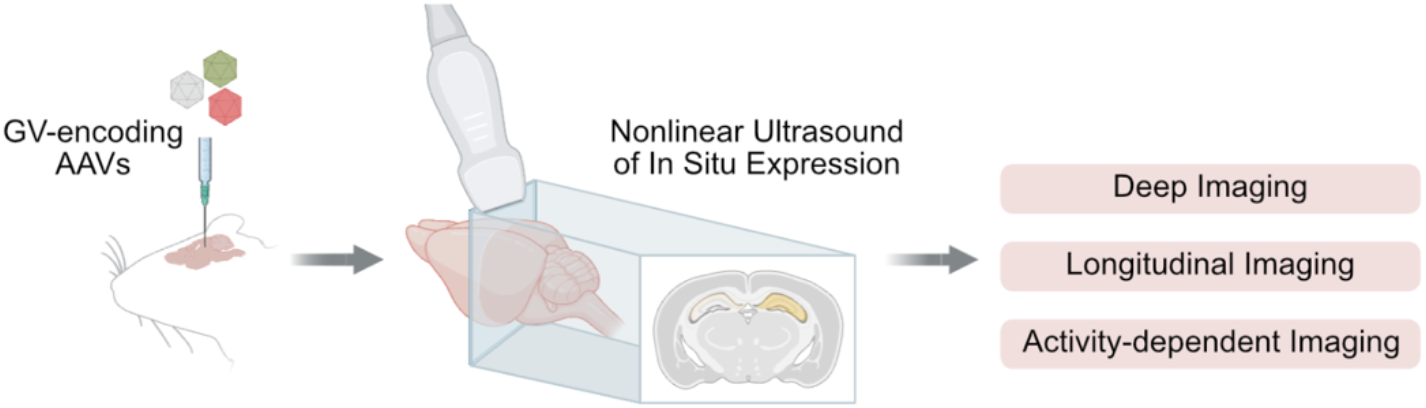
AAV-mediated delivery of GV genes into tissues enables ultrasound imaging of endogenous cellular function. Schematic showing injecting a 3-vector AAV system encoding the GV genes into the brain, followed by nonlinear ultrasound imaging of in situ gene expression and downstream capabilities, including tracking activity-dependent gene expression.

## RESULTS

### A three-vector AAV system enables robust GV expression in neurons

AAVs represent the most common viral delivery system used to deliver genetic payloads into tissues such as the brain, offering cell-type specificity and tissue tropism through engineered serotypes^8–12^. Therefore, we selected AAV as the vehicle to deliver the GV gene cluster into primary cells. We initially validated our genetic constructs in cultured primary neurons, a non-dividing cell type that served as a prelude to in vivo experiments in the mouse brain. We chose this cell type due to its common targeting with AAVs and the high interest in imaging its gene expression in the brain.

Inspired by previous work on GV expression in ex vivo modified immune cells^5^, we designed a three-vector AAV system expressing GVs under the control of a Tet transactivator, with each vector adhering to the ∼4.8 kb packaging size limit (**Fig. 2a**). The smallest GV gene cluster previously shown to express functional GVs is derived from *Anabaena flos-aquae*, with its assembly factor genes (*gvpNJKFGWV*) comprising 4.7 kb when linked by 2A peptides for polycistronic expression^4^. To accommodate the addition of promoters and polyA sequences, we split the assembly factor genes across two AAVs: one encoding *gvpNJK* and the transactivator rtTA (4.5 kb, ITR-ITR), and the other encoding TRE-driven *gvpFGWV* as well as GFP for quantifying transduction efficiency (4.2 kb, ITR-ITR). We placed the main structural GV gene, *gvpA*, co-expressed with RFP on a third AAV (2.6 kb, ITR-ITR), enabling independent control of its titer relative to the assembly factors. Each vector incorporated an engineered compact WPRE and poly-A sequence, CW3SL^13^, which occupies only a 432 bp DNA footprint, thereby minimizing packaging size. We packaged these vectors in AAV9, a serotype commonly used in neurons, and known for its global expression in the brain^14^.

**Figure 2:**
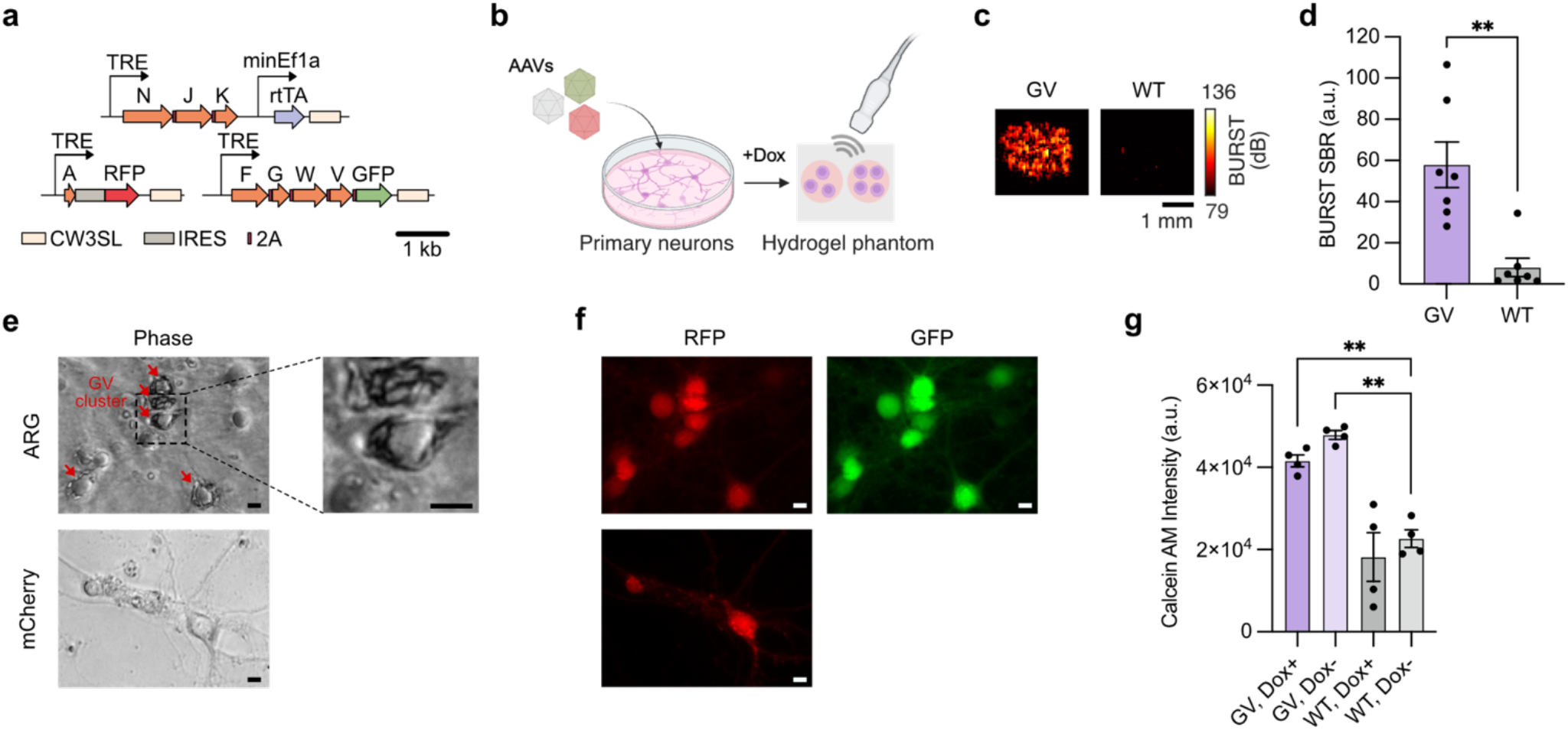
AAV-mediated delivery of the GV gene cluster into primary neurons. **a**, Schematic of the 3-vector AAV system, with the structural gene *gvpA* encoded on a separate plasmid to enable stoichiometric tuning of its expression relative to assembly factors. **b**, Schematic of the experimental protocol for validating GV expression in cultured mouse embryonic neurons. Cultured neurons were induced with doxycycline (Dox) every 24 hours, for 72 hours prior to dissociation and ultrasound imaging in agarose phantoms. **c**, Representative BURST images of neurons expressing GVs (left) and untransduced wild-type (WT) neurons (right). **d**, Quantification of the BURST signal-to-background ratio (SBR) in wells containing GV-expressing neurons compared to WT cells. Error bars represent mean ± s.e.m., *n* = 7 biological replicates. Each data point is the arithmetic mean of *n* = 2 technical replicates. Welch’s two-tailed t test, p = 0.0031. **e**, Phase contrast images of neurons transduced with GV-expressing AAVs or mCherry-expressing AAVs. Arrows point at visible GV clusters, appearing as dark structures inside the cell, under phase contrast microscopy. (Right) Magnified view of the cells outlined in the box, showing visible GV clusters. **f**, Corresponding fluorescence images of the neurons in (e). **g**, Calcein AM intensity in cultured neurons transduced with GV-expressing AAVs or untransduced WT cells, either induced with Dox (Dox^+^), or left uninduced (Dox^−^). n = 4 biological replicates. One-way ANOVA test with multiple comparisons correction. P-values from left to right: 0.0037, 0.002. Significance levels: n.s. p ≥ 0.5, *p < 0.05, **p < 0.01, ***p < 0.001.

To test this system, we cultured mouse cortical and hippocampal neurons in vitro and delivered the GV gene cluster using a 5:1:1 ratio of the AAV vectors encoding *gvpA* to those encoding the two sets of assembly factors (AFs). This ratio was chosen based on prior findings that an elevated *gvpA* ratio enhances the nonlinear acoustic scattering of GVs^4,5^. Expression was induced by adding doxycycline (Dox) every 24 hours for 72 hours. Cells were then trypsinized and embedded in agarose phantoms for ultrasound imaging (**Fig. 2b**).

BURST, a highly sensitive nonlinear detection method for GV imaging^15^, revealed a strong signal of 55.06 ± 14.59 a.u. (mean ± s.e.m.) in wells containing GV-expressing neurons, compared with 1.66 ± 0.06 a.u. (mean ± s.e.m.) in wells containing untransduced wild-type (WT) neurons (**Fig. 2c-d**). Phase microscopy of the cultures showed clusters of GVs^16,17^ forming in the soma (**Fig. 2e-f**). These clusters were absent in both untransduced cells and in cultures transduced with a constitutively expressing mCherry virus (**Extended Data Fig. 1a-c**). Calcein AM staining of Dox-induced and Dox-uninduced cells showed similar levels of metabolism across these two conditions in both AAV-transduced and untransduced WT cells, while the metabolic activity was generally higher in AAV-transduced cells, potentially reflecting a general cellular response to viral transduction (**Fig. 2g**).

These results demonstrate that a multi-AAV approach can successfully deliver ARGs to non-dividing primary cells, leading them to robustly express GVs visible with ultrasound and phase microscopy.

### AAV delivery of GV genes enables longitudinal deep-brain imaging of in situ gene expression

Having validated our multi-AAV expression system in vitro, we set out to test its ability to deliver the GV gene cluster to native cells inside an opaque tissue and enable longitudinal imaging of gene expression in a living animal over an extended period. The brain presents an ideal testing ground for this purpose, with established interest in using gene expression to visualize anatomical structures, cell types and activation. As our anatomical target, we selected the hippocampus, a tissue with a well-defined three-dimensional geometry, a location that is too deep for external optical access, critical importance in cognitive function and memory encoding, and malfunction associated with diseases such as epilepsy^18,19^.

We co-injected a mix of the three GV-encoding AAVs at a 5:1:1 ratio of *gvpA* to the assembly factor vectors into the right hippocampus of immunocompetent C57BL/6J mice. After 3-6 weeks, we induced GV expression for three days via daily intraperitoneal (IP) injections of Dox, then imaged the brain through a cranial window, which eliminates acoustic scattering from the skull and maximizes GV-specific signal (**Fig. 3a**). For comparison, we also injected mice with an AAV virus constitutively expressing mCherry at similar titers and stereotaxic coordinates. Imaging with a linear array ultrasound transducer along the anterior-posterior axis, we acquired BURST images of GVs and Doppler images of the vasculature as anatomical reference. We observed strong BURST signal in the hippocampus of mice injected with the GV-expressing AAVs at 4.0e4 ± 1.5e4 a.u. (mean ± s.e.m.), while mCherry-injected controls exhibited minimal signal at 3.8e2 ± 2.4e2 a.u. (mean ± s.e.m.) (**Fig. 3b, d** and **Extended Data Fig. 2**). Interestingly, despite injecting the vectors unilaterally, we also observed BURST signal in the contralateral hippocampus, demonstrating the ability of this approach to visualize gene expression even at distant projection sites, which are due to contralateral projections from the ipsilateral hippocampus to the opposite side across the midline^20^. The ultrasound-observed expression pattern matches what we saw by post-mortem fluorescence in the same animals (**Fig. 3c**).

**Figure 3:**
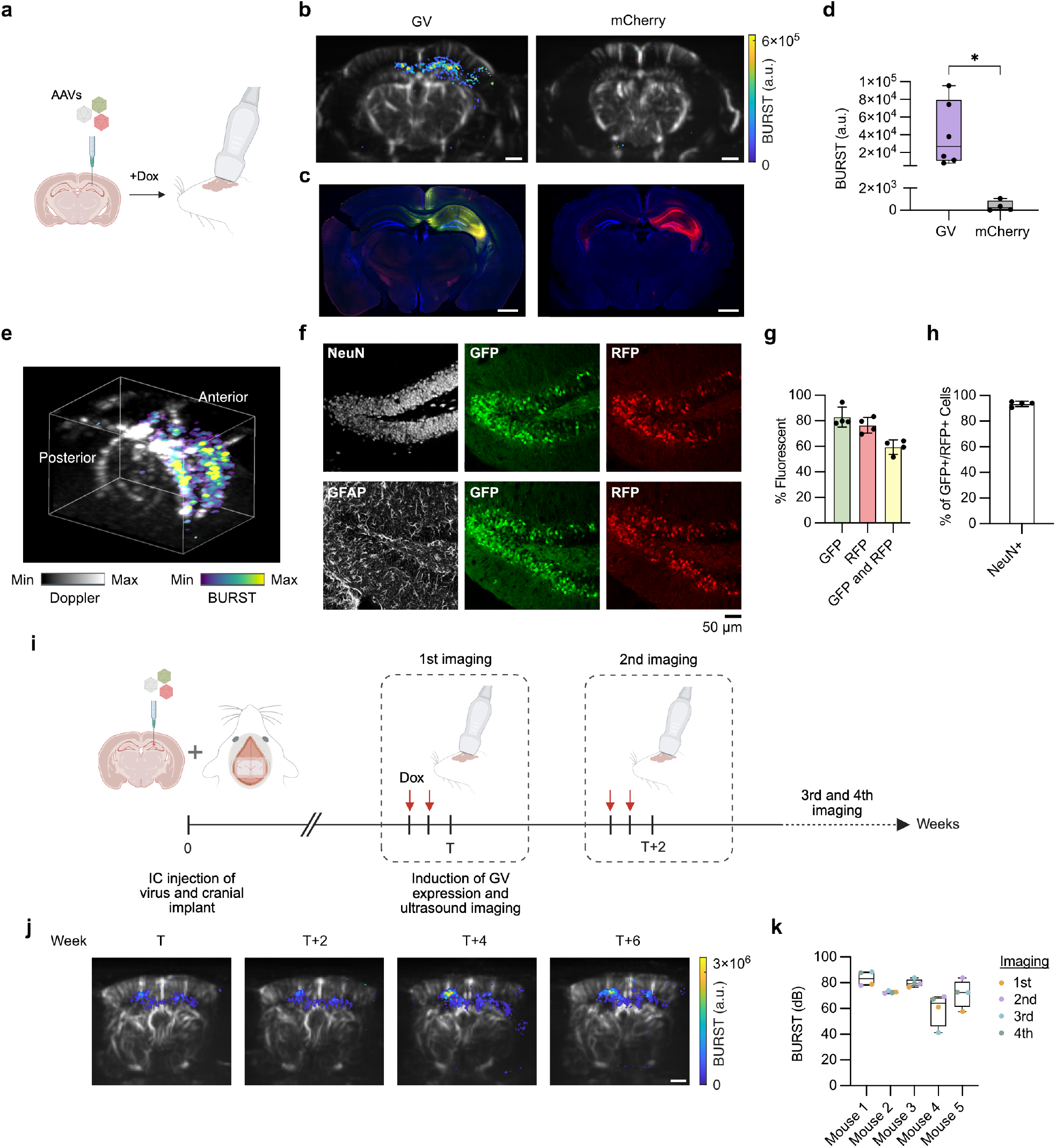
AAV delivery and longitudinal imaging of GV-expression in the mouse brain. **a**, Schematic of the experimental approach. A mix of three GV-encoding AAVs is injected intracranially into the hippocampus. GV expression is induced for 72 hours, and the hippocampus is imaged through a cranial window. **b**, Representative BURST images (colormap) overlaid on Doppler images (grayscale) of mice injected with GV-encoding AAVs or a constitutively expressing mCherry AAV as control. **c**, Corresponding histological sections. Blue: DAPI, Green: GFP, Red: RFP in the left image and mCherry in the right image. **d**, Quantification of BURST signal in mice injected with GV AAVs (n = 6) versus mCherry AAVs (n = 4). Welch’s one-tailed t test, p = 0.0203. **e**, Volumetric BURST imaging of the brain using Takoyaki imaging, overlaid on volumetric Doppler image. Anterior and posterior directions of the brain are indicated. **f**, Representative histological images from the dentate gyrus (DG) showing GFP and RFP expression, co-stained with NeuN (neurons) and GFAP (astrocytes). **g**, Quantification of fluorescent cell populations in the DG, showing percentages of GFP^+^, RFP^+^, and GFP^+^/RFP^+^ cells. n = 4 mice. n = 417 ± 106 (mean ± s.d.) cells counted per mouse. **h**, Fraction of GFP^+^/RFP^+^ cells that colocalize with NeuN. n = 4 mice. n = 252 ± 81 (mean ± s.d.) cells counted per mouse. **i**, Schematic of the longitudinal imaging experiment. Mice were injected with the AAV cocktail, and an acoustically transparent cranial window was implanted. GV expression was induced for 48 hours prior to each imaging session, followed by ultrasound imaging and GV collapse. This cycle was repeated every 2-3 weeks for four rounds. IC: intracranial. **j**, Representative ultrasound images of the same anterior-posterior plane (within 0.5 mm accuracy), acquired over 6 weeks across four imaging sessions. **k**, Quantification of BURST signal in the same imaging plane across four sessions in n = 5 mice. In (d) and (k), the line in the middle represents the median, the box spans the 25th to 75th percentiles, and the whiskers indicate the minimum and maximum values. In (g) and (h), the line represents the mean and error bars indicate the s.e.m. Scale bars: 1 mm, unless noted otherwise.

BURST imaging with a linear array enabled high-resolution visualization of GV expression in individual 2D coronal planes of the brain. However, each BURST acquisition can collapse GVs in adjacent elevation planes and peri-focal axial regions without capturing their nonlinear signal, making it challenging to capture GV expression in the entire brain with a mechanically sweeping linear array. To demonstrate that our reporter genes are compatible with larger-coverage volumetric imaging, we performed Takoyaki 3D BURST imaging using a matrix array probe^21^. Takoyaki imaging scans its underlying 3D volume by transmitting multiple focused beams simultaneously along both the lateral and elevational axes, with more axially homogeneous pressure fields. Using this method, we captured nonlinear ultrasound signal across the entire hippocampus—from anterior to posterior—in a single acquisition (**Fig. 3e** and **Supplementary Video 1**).

In post-mortem histological quantification, we found that, among fluorescent cells, 59.4 ± 2.8% (mean ± s.e.m., n = 4 mice) were both GFP and RFP positive, indicating successful triple transduction with the three GV-encoding AAV vectors (**Fig. 3f, g**). Within this triple-transduced population, 93.5 ± 1.0% of the cells (mean ± s.e.m.) co-localized with NeuN, a neuronal marker, confirming that the majority of BURST signal originated from neurons (**Fig. 3f, h**). Moreover, 87.9 ± 2.5% (mean ± s.e.m.) of the triple transduced cells were negative for cleaved caspase-3, an apoptosis marker (**Extended Data Fig. 3**), confirming that the vast majority of GV-expressing cells are viable. The small percentage of cells that were caspase-positive may be apoptotic due to high levels of viral titer or excessive GV gene expression.

Having established robust in situ GV expression and imaging, we next tested whether our system could support longitudinal imaging. In a separate set of animals, we injected the GV-expressing AAV mix into the right hippocampus and implanted an acoustically transparent cranial window for repeated high-sensitivity imaging^22,23^. In each of four sessions separated by 2-3 weeks, we induced GV expression for 48 hours, performed BURST imaging, and collapsed all GVs using ultrasound (**Fig. 3i**). While some variability in expression was observed between imaging sessions, we saw clear BURST signal in each session without evidence of signal loss over time (**Fig. 3j, k and Extended Data Fig. 4**).

Together, these results establish the feasibility of using ultrasound to image in situ gene expression in opaque, intact living organisms across time.

### GV expression driven by IEGs tracks intracellular activity

We next sought to connect GV expression to cellular function. Gene constructs in which fluorescent protein expression is driven by IEGs have been widely used to visualize spatial patterns of brain-wide neural activity^24–26^. However, as a post-mortem histological readout, this approach is inherently limited to terminal experiments. If IEG activity could be visualized with ultrasound instead of light, it would enable a non-terminal and dynamic readout of activity patterns.

To develop an IEG-driven acoustic reporter construct, we focused on two IEG promoters: the widely used *cFos* promoter and RAM, a synthetic promoter with enhanced expression fold change that combines a synthetic enhancer module consisting of Activator Protein 1 (AP-1) and Npas4 sites with a minimal *cFos* promoter^26^. To minimize leakiness and allow more precise temporal control over GV expression, we reduced the half-life of the rtTA transactivator by adding the mouse ornithine decarboxylase (ODC) degron^26,27^ to its 3’ end with a GS linker, creating a variant we termed “RAM-deg”. In our GV expression constructs, we replaced the constitutive minEf1a promoter upstream of rtTA with these activity-inducible promoters and tested them in HEK cells.

We incubated the cells with an activity induction cocktail, containing phorbol 12-myristate 13-acetate (PMA) and ionomycin, and Dox for 4-6 hours (**Fig. 4a**). 18-20 hours later, we observed significant BURST signal from all three promoters. Among them, the RAM-deg promoter produced the highest fold change in BURST signal, reaching 5.6× compared to resting state (**Fig. 4b, c**). Flow cytometry confirmed that RAM-deg had the lowest basal activity and the highest fluorescence expression upon induction (**Fig. 4d**). Based on these findings, we selected the RAM-deg construct for further experiments.

**Figure 4:**
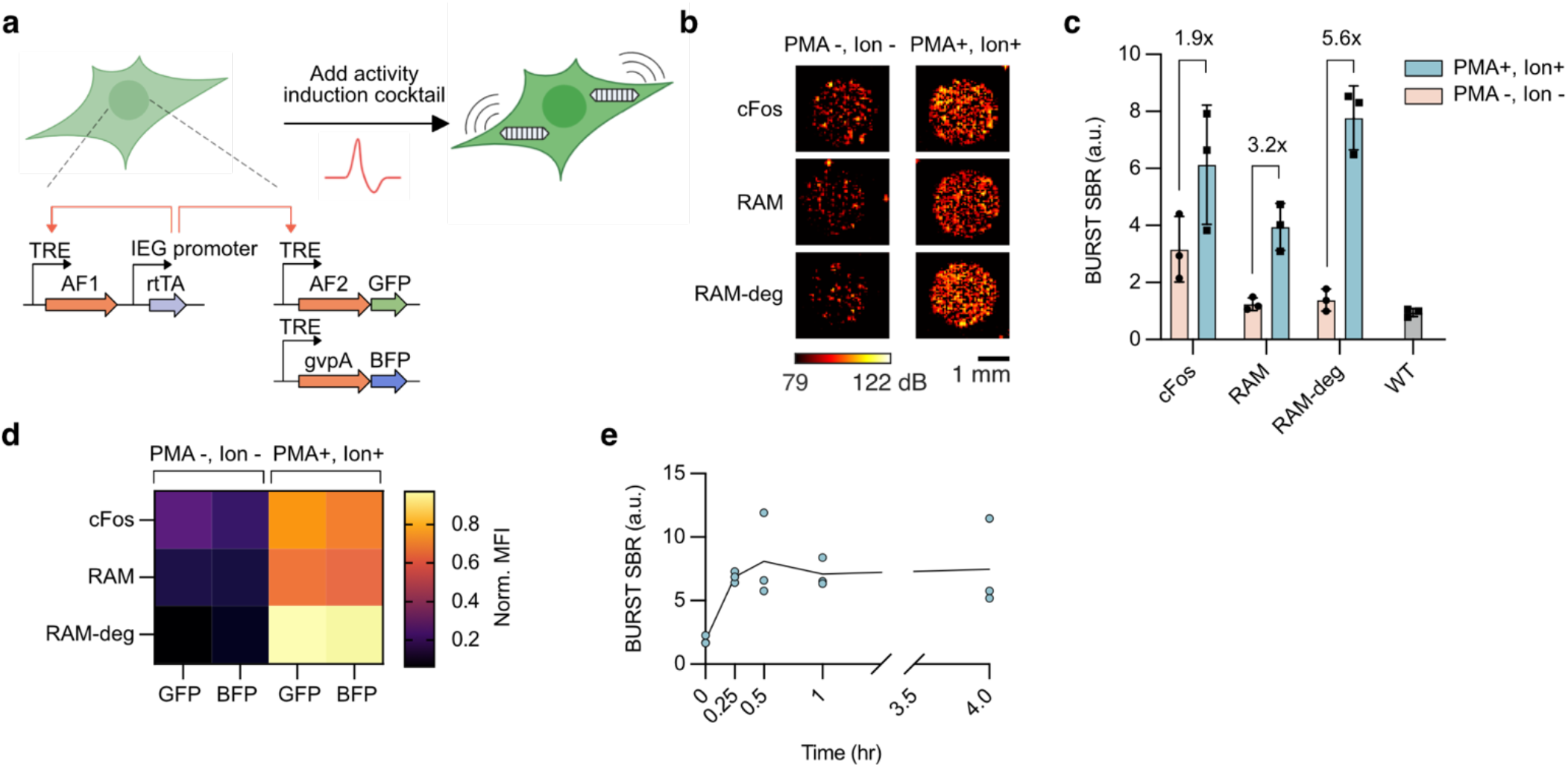
IEG promoters enable activity-dependent GV expression in HEK293T cells. **a**, Schematic of lentiviral plasmid design and experimental approach for testing variants of IEG-induced GVs in HEK293T cells induced with an activity induction cocktail (PMA and ionomycin). rtTA is expressed under an IEG promoter (either cFos or RAM). To reduce the half-life of rtTA, a degron-linked version (RAM-deg) was also tested. AF1: assembly factor cassette 1 (*gvpNJKF*); AF2: assembly factor cassette 2 (*gvpGWV*). Schematic of genetic elements does not represent their relative sizes. **b**, Representative BURST images of HEK293T cells expressing IEG-induced GV variants in their resting (PMA-, Ion -) or active (PMA+, Ion +) state. **c**, Quantification of BURST signal and its fold change in cells expressing different IEG variants compared to untransduced WT cells. **d**, Mean fluorescence intensity (MFI) of resting and activity-induced cells expressing different IEG variants, normalized to the maximum intensity for each fluorescence channel. **e**, BURST signal in RAM-deg-expressing cells incubated with the activity induction cocktail for varying durations of time. Error bars represent mean ± s.e.m. n = 3 biological replicates. Each data point represents the arithmetic mean of n = 2 technical replicates. Ion: ionomycin.

To evaluate the sensitivity of the RAM-deg GV expression system, we tested how its response changed with the duration of activity induction. AP-1-mediated activation of the *cFos* promoter is transient—peaking within 0.5-2 hours of activation and decaying by 4-6 hours^28,29^. Therefore, we stimulated cells for varying durations of time, from 15 minutes to 4 hours. We found that even 15 minutes of stimulation was sufficient to produce detectable GV expression (**Fig. 4e**).

These results establish RAM-deg as a sensitive, activity-inducible system for tracking IEG-induced transcription with ultrasound.

### Ultrasound imaging of conditional IEG-induced gene expression in the brain resulting from epileptic seizures

One of the main applications of post-mortem immunohistochemistry of IEGs has been to visualize spatial patterns of neural activity in response to various stimuli, behaviors or disease states^30–36^. However, conventional histology approaches to visualize IEGs require sacrificing the animal, making it impossible to visualize activity in individual subjects over sequential timepoints and increasing the use of animals in studies. We therefore set out to demonstrate the use of ARGs to perform longitudinal and non-invasive imaging of IEG transcription in a brain activation paradigm.

For this first-of-its-kind demonstration, we used a well-characterized model of temporal lobe epilepsy in which systemic kainic acid injections induce seizures^37,38^. This results in strong activation of the hippocampus, and is a well-established paradigm to validate IEG-driven reporters^26^. We confirmed that, in our hands, kainic acid reliably increases *cFos* expression in the hippocampus via immunohistochemistry (**Extended Data Fig. 5**).

To enable in vivo IEG imaging, we adapted the RAM-deg architecture into our AAV GV expression backbone by replacing the minEf1a-rtTA element (from **Fig. 2a**) with RAM-deg, the destabilized rtTA expressed under the RAM promoter. We also added a constitutively expressed iRFP670 fluorophore^39^ as a marker of transduction, confirming in a separate experiment that its addition did not affect in vivo GV expression (**Extended Data Fig. 6**).

To image seizure-induced changes in gene expression, we injected RAM-deg AAVs into the right hippocampus and implanted an acoustically transparent window on the brain. We divided the mice into two groups, an experimental group with induced seizures, and a control group that received sham treatment. We performed two sequential imaging sessions (baseline and after seizure/sham) on both groups. For the first session (baseline), we administered Dox and saline to both groups, followed by ultrasound imaging 24 hours later (day 1). For the second session that was performed 6 days later, we induced seizures in the experimental group via IP injections of kainic acid together with Dox, while the control group received a second dose of saline and Dox as sham treatment (**Fig. 5b**). Both sets of mice were imaged 24 hours later (day 7).

**Figure 5:**
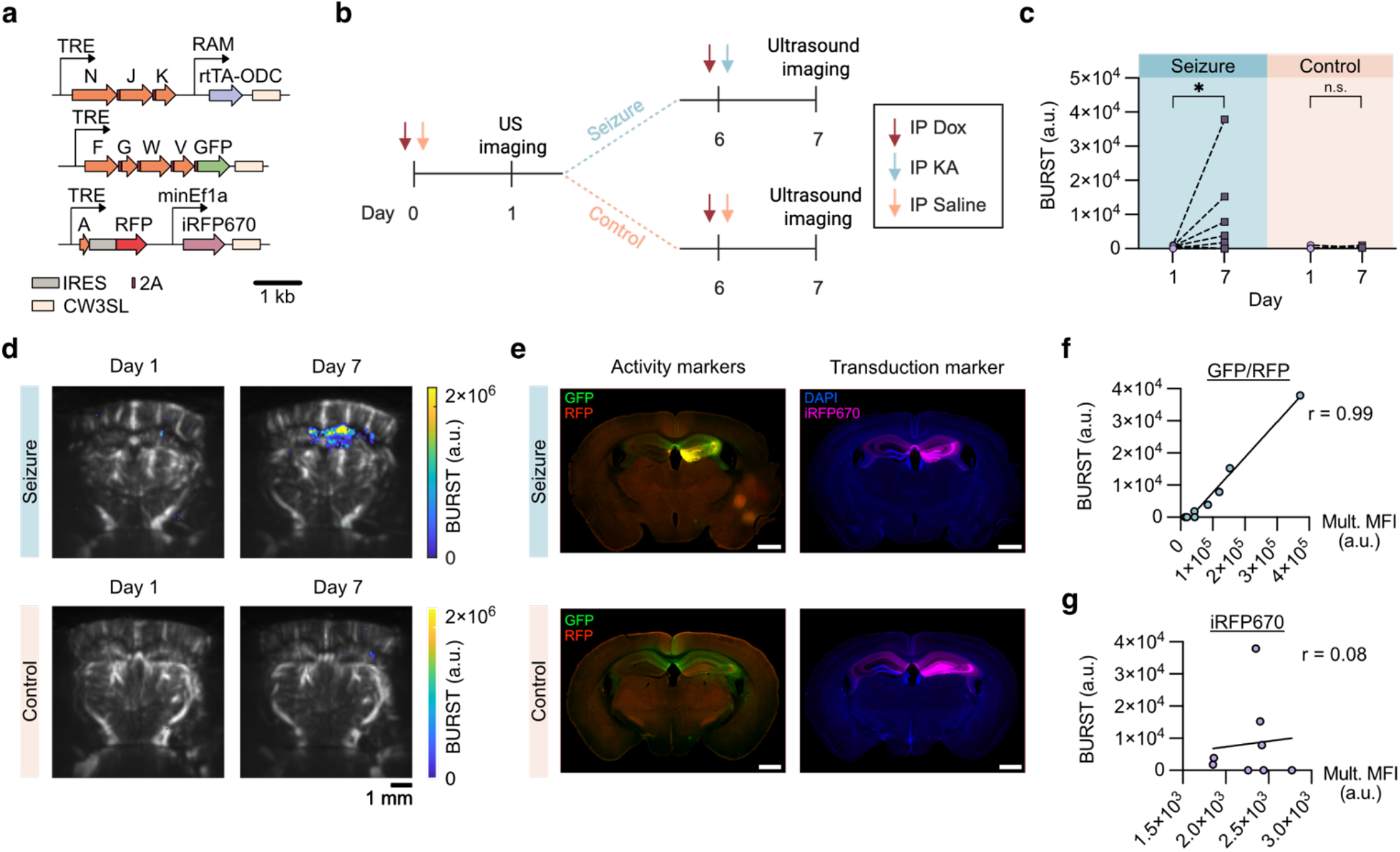
IEG-induced GV expression tracks changes in cellular activity in the hippocampus caused by seizures. **a**, Schematic of the AAV design for IEG-induced GV expression using the RAM-deg promoter variant. A constitutive fluorophore, iRFP670, was included on the *gvpA* construct to account for variability in transduction efficiency. **b**, Experimental timeline for seizure induction and activity imaging. Mice were imaged at two time points: day 1 (baseline), following IP injections of Dox and saline; and day 7, following either kainic acid (KA) and Dox (seizure group) or a second saline and Dox (control group) injection. **c**, Quantification of BURST signal in the hippocampus of seizure-induced (n = 8 mice) and control (n = 4 mice) groups, as described in (b). Each line represents an individual mouse. Seizure group mice that did not exhibit an increase in fluorescence despite receiving KA were excluded from the significance test. Ratio-paired t test, p (Seizure) = 0.003, p (Control) = 0.284. Significance levels: *p ≤ 0.5, n.s. p > 0.5. **d**, Representative ultrasound images from seizure-induced mice showing activity-induced BURST signal in the hippocampus and control mice. **e**, Fluorescence images of histological sections corresponding to the mice shown in (d), acquired at matched anterior–posterior planes (±0.3 mm). Left column: GFP and RFP fluorescence driven by the IEG promoter. Right column: constitutive iRFP670 expression and DAPI staining. **f**, Correlation between the BURST signal and fluorescence signal in the corresponding histological sections from the seizure group. Multiplied MFI (Mult. MFI) is defined as the product of GFP and RFP channel mean fluorescence intensities in the right hippocampus normalized to the contralateral side. Pearson r = 0.992, p (two-tailed) < 0.0001, n = 8 mice. **g**, constitutive iRFP670 fluorescence from histological sections. Pearson r = 0.082, p (two-tailed) = 0.846, n = 8 mice.

In the seizure group, we observed a large increase in ultrasound signal in the hippocampus on day 7 compared to baseline measurements taken on day 1 (**Fig. 5c, d**). In contrast, we observed no significant change in ultrasound signal in the control group on day 7 versus day 1 (**Fig. 5c, d**). Histological analysis supported these findings, revealing strong GFP and RFP fluorescence in the hippocampus of seizure-induced mice, and weak fluorescence in controls (**Fig. 5e**).

Since we observed some heterogeneity in BURST signal intensity among the seizure group (**Fig. 5c** and **Extended Data Fig. 7a, b**), we wondered whether this variability was due to inconsistencies in ultrasound imaging or if it actually represented underlying expression patterns. Histological analysis showed that mice with low BURST signal also had weaker GFP and RFP fluorescence, comparable to fluorescence expression in the control group (**Extended Data Fig. 7c**). In fact, across the seizure group, we observed a very strong correlation between BURST signal and the fluorescence intensity of GFP and RFP in the hippocampus (**Fig. 5f**; Pearson r = 0.992, p < 0.0001). In contrast, there was no significant correlation between BURST signal and the constitutive fluorophore iRFP670 (**Fig. 5g**; Pearson r = 0.082, p = 0.846).

These experiments establish the ability of ARGs to visualize physiological changes in gene expression in vivo in a way that is quantitatively consistent with conventional post-mortem fluorescent readouts. The major difference is that animals imaged with ultrasound remain alive and can be imaged repeatedly at multiple timepoints (as shown here on day 1 and day 7). By eliminating the need for terminal histology and enabling repeated measurements in the same subject, this platform represents a major advancement for studying tissue-wide circuit dynamics over time.

## DISCUSSION

Imaging gene expression directly within living organisms provides a powerful window into the dynamics of in vivo cellular function, but has remained technically challenging beyond superficial tissues or ex vivo models. This work introduces an approach to overcome this limitation by, for the first time, expressing GVs in situ in a live animal and enabling ultrasound-based imaging of dynamic physiological gene expression in intact tissue. Using the brain as a testing ground, the multi-AAV system introduced here enables robust expression of GVs following intracranial injections, resulting in strong ultrasound contrast in the hippocampus. This signal is detectable through optically opaque overlying tissue and can be captured in a single 3D acquisition. We demonstrate that this platform supports longitudinal imaging in the same animal over multiple weeks.

Furthermore, we developed a minimally leaky activity-dependent acoustic reporter system regulated by endogenous transcription factors, that sensitively tracks changes in intracellular activity in both in vitro and in vivo models. We demonstrated the use of this reporter in imaging IEG expression in the hippocampus after epileptic seizures. Leveraging the deep penetration and non-invasiveness of ultrasound, this tool enables paired imaging of baseline and post-stimulation gene expression in the same animal, and helps to capture variability in activity patterns caused by seizures in different subjects^40^.

Future work could extend this imaging platform to brain-wide monitoring of gene expression following systemic AAV delivery and the use of non-invasive IEG imaging to investigate other dynamic processes such as memory encoding within the same animal over extended time periods^33,41–44^. When paired with systemic delivery methods or transgenic rodent models, this technique has the potential to support longitudinal, multi-organ circuit analysis with cell type specificity. Furthermore, integrating our AAV expression system with recently developed GV-based protease^6^ or calcium sensors^7^ could pave the way for non-invasive imaging of spontaneous in situ molecular dynamics with high spatial and temporal resolution.

Although we did not observe any gross behavior changes or significant cytotoxicity in animals expressing GVs, further work is needed to assess the impact of GV expression on cellular physiology, ideally in a cell-type–specific manner using functional assays. Another open question is whether co-transduction efficiency of all three AAVs could become a limiting factor in the context of systemic delivery. Reducing the system to two AAVs, which could be achieved through rational engineering or further screening of more compact GV gene clusters, could improve efficiency and versatility^4,45^. While we demonstrated that ultrasound could image GV expression across large brain regions, future studies should explore the minimum viral dose required to resolve smaller substructures. Future studies are also needed to examine GV expression efficiency in additional cell types in the brain and other organs. Lastly, improving the nonlinear acoustic contrast of GVs through the development of engineered variants or the tuning of gene stoichiometry could further enhance imaging sensitivity and allow the use of non-collapse-based methods of imaging^46,47^.

Altogether, this platform establishes the first genetic toolset for deep-tissue, ultrasound-based imaging of in situ gene expression in live animals. By enabling longitudinal, non-invasive visualization of transcriptional dynamics, it paves the way for studying cellular function over time within intact tissues and in natural physiological contexts.

## METHODS

### Plasmid construction and molecular biology

All genetic constructs were made using either KLD mutagenesis or Gibson Assembly with enzymes from New England BioLabs. The plasmids and their corresponding genetic material sources are detailed in Supplementary Table 1. Constructs were cloned into NEB Turbo Competent E. coli (High Efficiency) (New England BioLabs, C2984). Fluorescent reporters referred to as GFP, RFP, and iRFP670 in the text correspond to mEGFP, mScarlet-I, and emiRFP670 respectively. All AAV transgenes were packaged into AAV9 plasmids, and both packaging and titering were outsourced.

### Dissection of embryonic mouse neurons

Tissue culture-treated 24-well polystyrene plates were coated with poly-D-lysine (PDL, 100 µg/mL) (Gibco, A3890401) diluted to 50 µg/mL in sterile DPBS, at room temperature for 1 hour. Poly-D-lysine solution was then removed, washed twice with molecular biology-grade water and left to dry prior to seeding.

Pups were extracted from a pregnant female C57bl/6J mouse (Charles River, strain #027) on embryonic day 18 (E18, Charles River Laboratories) under a humane protocol approved by the Institutional Animal Care and Use Committee of the California Institute of Technology. They were placed on ice while still enclosed in membranous sacs. Under a sterile hood brains of pups were extracted one at a time and immediately placed in cold Hank’s Balanced Salt Solution (HBSS). The cortex and hippocampus from each hemisphere were dissected and stored in cold HBSS. Dissected tissue was then cut into small pieces. For enzymatic dissociation, tissue fragments were incubated in papain (Worthington Biochemicals PAP2/LK003178, diluted to 15– 20 U/mL in 1X PBS) for 20–30 minutes at 37 °C, with gentle shaking. Digestion was stopped by transferring tissue into 10% FBS in HBSS, while minimizing shear stress. Cells were then gently triturated. The suspension was centrifuged at 100 × g for 8 minutes. The resulting pellet, containing a homogeneous suspension of single cells, was resuspended in neuronal culture media containing Neurobasal Plus media (Thermofischer, #A3582901), 1X B27 Plus supplement (Thermofischer, #A3653401), and 1X penicillin/streptomycin. Cells were counted with AO/PI dye. 2.5 × 10^5^ cells were plated in each well of a pre-coated 24 well plate. The culture media was fully replaced with pre-warmed media on day in vitro 1 (DIV 1), followed by half-media changes every four days thereafter.

### Transduction and induction of primary cultured neuron

Cultured neurons were transduced on DIV 5 with either 1.4 × 10^6^ GC/cell of the three GV-encoding AAV9 viruses or 1 × 10^6^ GC/cell of a single mCherry-encoding AAV9 virus (Addgene, #114470). Beginning on DIV 13, neurons were induced every 24 hours with Dox (1 µg/mL). Imaging was performed on DIV 16.

On DIV 17, Calcein AM (AAT Bioquest, #22012) staining was performed according to manufacturer’s protocol. Cells were once washed in serum-free media, and incubated with Calcein AM (4 µM), for 30 min at 37°C and 5% CO2. Cells were then washed once with serum-free media, and imaged with the Tecan Spark plate reader. Signal from un-stained wells was used for background subtraction.

### In vitro IEG induction of HEK239T cells

HEK293T cells were transduced with the activity-dependent three vector lentiviral system, using previously established methods for transduction of lentivirally-encoded GVs^5^, at MOI 9:3:3 of *gvpA*-encoding virus to the two assembly factor viruses. Cells were seeded ∼24 hours before induction in DMEM (Corning, 10-013-CV) with 10% FBS (Gibco) and 1X penicillin/streptomycin at 37 °C, 5% CO2, until they reached 80-90% confluence. They were then incubated in ionomycin (1.3 µM) (Sigma, #I9657), phorbol myristate acetate (PMA, 100 ng/mL) (VWR, Catalog 16561-29-8), and Dox (1 µg/mL) supplemented DMEM at 37 °C, 5% CO2 for 4 hours. The inducers were removed, and the cells were washed once in 1X PBS and maintained in Dox (1 µg/mL) supplemented DMEM media for 20 hours before ultrasound imaging and flow cytometry with the MACSQuant Analyzer 10 (Miltenyi Biotec). In time induction experiments, the duration of induction was varied, as shown in **Fig. 3e**.

### Phantom preparation for in vitro ultrasound imaging

On the day of imaging, cells were trypsinized with trypsin (0.25% for HEK cells, and 2.5% for cultured neurons) for 5 minutes at 37°C. They were then gently detached from the culture dish and resuspended in 1X PBS. DNase I (100 µg/mL) was added to the suspension containing cultured neurons to prevent clumping. The cell suspension was then centrifuged at 300 × g for 5 minutes, and the supernatant was carefully removed. Cells were then briefly mixed at a 1:1 ratio with 1% melted agarose in 1X PBS at 42°C. The agarose solution was prepared at least 24 hours in advance to eliminate bubbles. The cell-agarose mixture was then loaded into agarose phantoms, yielding a final cell concentration of approximately 25 million cells/mL for HEK293T cells and 6 million cells/mL for neurons.

### Animals

C57BL/6J mice were obtained from Jackson Lab and bred at Caltech facilities (strain #000664). Animals were housed in a 12 h light/dark cycle and were provided with water and food ad libitum. A mixed population of male and female mice, chosen randomly, were used in studies. All experiments were conducted under a protocol approved by the Institutional Animal Care and Use Committee of the California Institute of Technology.

### Intracranial injections

Solutions of the three GV-encoding AAV9 viruses, mixed at a 5:1:1 ratio of the *gvpA*-encoding virus to the assembly factor viruses, or a single AAV9 expressing mCherry under the Ef1a promoter were injected into the hippocampus of C57BL/6J mice at 8-11 weeks of age using a pulled glass needle coupled with a motorized pump at 70 nL/min using a stereotaxic frame. All injections were done at coordinates of -2.1 mm anterior-posterior (AP), -1.5 mm medio-lateral (ML), -1.8 mm dorso-ventral (DV) with respect to bregma. The needle remained in place after injection for 5 min to avoid backflow along the needle tract. The total volume of injection was 3 µL, for a total viral load of 3 × 10^10^ to 3 × 10^11^ genome copy (GC) number, measured by qPCR titer. For each batch of virus, an initial test experiment was performed to determine the amount of time needed to wait after injection before a significant BURST signal was detected.

### Cranial window surgery

Ultrasound imaging of mice that were scanned only once was performed after skull removal. Specifically, mice were anesthetized under 1-2.5% isoflurane and injected subcutaneously with mannitol to reduce brain swelling. A craniotomy was performed from AP - 0.5 mm to AP -3.5 mm and from ML -2.5 mm to ML +2.5 mm using a micro-drill steel burr (Fine Science Tools).

### Acoustically transparent cranial window implants

To allow longitudinal imaging, the skull of mice was partially removed and replaced with an acoustically transparent window. Specifically, the skull flap was replaced with a TPX (polymethylpentene) (Goodfellow #692-976-87) window of similar size and fixed in place using a UV-curable composite (Tetric EvoFlow, Ivoclar) and dental cement (C&B Metabond, Parkell). The TPX implant surgery and the ultrasound imaging were spaced at least one week apart to ensure there were no bubbles under the TPX window.

### Doxycycline administration

For single-time imaging experiments, mice received IP injections of doxycycline hydrochloride (Dox HCl) at 4 mg/kg in saline every 24 hours for 72 hours. For repeated imaging experiments, mice were injected with Dox HCl every 24 hours for 48 hours prior to imaging.

### Seizure inductions

All mice used for seizure induction, including controls, were single-housed following intracranial injections. To ensure adaptation, mice were transferred to fresh housing with access to standard chow and water at least three days before behavioral experiments. They remained in the same cage throughout the study. All procedures were conducted during the light cycle, and careful handling was employed to minimize stress.

IP injections were performed under brief anesthesia with 3.5% isoflurane. To induce transgene expression, all mice first received an IP injection of Dox HCl (4 mg/kg in saline) and were returned to their housing. After a 30-minute interval, mice were re-anesthetized under 3.5% isoflurane and administered an IP injection of either saline (200–250 µL) or kainic acid (20 mg/kg in saline, pH 7.4) (Bio-Techne, #7065). In the seizure group, if no behavioral seizure symptoms were observed within one hour of kainic acid injection, an additional 5 mg/kg dose was administered.

Following kainic acid or saline injection, mice were continuously recorded for one hour using a webcam positioned above the cage to capture behavioral seizure activity. Animals that did not recover from seizures within two hours received a subcutaneous injection of diazepam (10 mg/kg in saline) and were closely monitored until full recovery. All mice were returned to their standard housing overnight and imaged 24 hours after Dox administration.

### Histological analysis

#### Tissue collection and cryosectioning

Following imaging, animals were perfused transcardially with 15 mL PBS and 15 mL 10% formalin. Brains were dissected, fixed in 10% formalin at 4 °C on a rotator for 12–24 h, then transferred to 30% sucrose for 48 h. Samples were embedded in OCT compound (Tissue-Tek) and stored at −80 °C. Frozen brains were sectioned coronally at 60 µm thickness (AP -1.2 mm to - 3.5 mm from bregma), collected in PBS, and stored at 4 °C in the dark for up to two weeks.

#### Immunohistochemistry

All steps were performed on an orbital shaker. Sections were washed once in PBS to remove OCT, then blocked for 60 min at room temperature. Standard blocking buffer contained 5% goat serum and 0.5% Triton X-100 in PBS (blocking for cleaved Caspase 3 (Casp3) used 10% donkey serum, 2% BSA, 0.5% Triton X-100). Sections were washed three times (5 min each) in PBS, incubated overnight at 4 °C with primary antibodies in 1% BSA, 0.3% Triton X-100 in PBS (or 10% donkey serum, 2% BSA, 0.3% Triton X-100 for Casp3). After three PBS washes, sections were incubated with secondary antibodies in PBS for 2 h at 4 °C in the dark, washed again, and, where indicated, stained with DAPI (2.5 µg/mL, 10 min). Sections were washed three final times in PBS, mounted with 200 µL mounting medium (Electron Microscopy Sciences, #17985-11), and sealed with a coverslip.

#### Antibodies

Primary antibodies: anti-NeuN (Abcam, ab104225, 1:1000), anti-GFAP (Cell Signaling Technology, #12389S, 1:250), anti-cleaved Caspase 3 (CST, #9661S, 1:400). Secondary antibodies: Alexa Fluor 647 (Abcam, ab150079, 1:1000), Alexa Fluor 405 (Abcam, ab175651, 1:1000).

#### Microscopy imaging

Whole-slice imaging was performed using a 10X objective on an epifluorescence microscope (Olympus VS120). Confocal imaging was conducted on an inverted laser scanning confocal microscope (20X objective) to acquire z-stacks (2 µm optical sections) of GFP, RFP, NeuN, and GFAP staining. For cleaved caspase-3 quantification, z-stacks were acquired on the same system using a 40X objective with 1 µm optical sections.

#### Image analysis and cell counting

Fluorescence images from the epifluorescence microscope were analyzed in QuPath. To quantify fluorescence intensity relative to BURST signal, a manual ROI was drawn around the hippocampus on the injected hemisphere to determine the MFI of GFP, RFP, or iRFP670. A background ROI was drawn contralaterally over the upper molecular layer of the dentate gyrus (DG) and the stratum lacunosum-moleculare (SLM) of CA1 (regions with minimal iRFP670 expression). Signal values were background-subtracted per sample.

Only confocal images were used for single cell counting. A 6 µm substack with strong expression/staining was selected in Fiji (ImageJ) using “Make Substack,” and each channel was exported in HDF5 format. Pixel segmentation was performed in Ilastik, then loaded into Fiji for analysis. Binary masks were generated from each segmentation. To restrict counts to nuclei, masks were intersected with DAPI using AND operation in the Image Calculator. Co-expressing cells were identified by logical AND operations between channels (e.g., GFP-DAPI and RFP-DAPI), followed by watershed segmentation to separate clustered objects. Cell counts were obtained using “Analyze Particles” with a minimum area threshold of 20 pixels.

For cleaved caspase-3 quantification, a Gaussian blur (σ = 0.4 µm) was applied in Fiji to reduce noise. Caspase-3-positive cells were manually counted using the Cell Counter plugin. Cell death was expressed as the fraction of cleaved caspase-3-positive among total GFP- and RFP-expressing cells.

### In vitro ultrasound imaging

#### *In vitro* BURST imaging sequence

Ultrasound imaging for in vitro experiments was performed using a Verasonics Vantage programmable ultrasound scanning system with an L22-14vX 128-element linear array transducer (Verasonics). Image acquisition was conducted using previously published BURST scripts^15^. The transmit waveform was set to a frequency of 15.6 MHz, with a 67% intra-pulse duty cycle and seven half-cycle transmits. The BURST pulse sequence consisted of an initial low-pressure frame (transducer voltage: 1.6 V; peak positive pressure: 0.4 MPa) followed by 45 high-pressure frames (transducer voltage: 15 V; peak positive pressure: 3.6 MPa). The programmable transmit focus was set to 8 mm to align with the fixed elevation focus of the transducer. All images were captured in AM mode.

#### *In vitro* BURST image processing

All in vitro BURST images were reconstructed using a previously established temporal-template unmixing algorithm across individual pixel locations in the frame stack^15^. Normalized signal-to-background ratio (SBR) was defined as the average BURST signal in the well containing a sample of interest divided by the average signal inside a noise ROI drawn adjacent to the well and inside the phantom.

In all images where decibel (dB) units are used, dB is defined as:

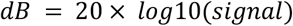

### In vivo 2D image acquisition with linear array

For all images captured using the linear array, an initial low-voltage power Doppler image (5V) was acquired to find the desired imaging plane. This voltage was selected based on preliminary experiments to prevent GV collapse. Once the imaging plane was established, a BURST image was acquired, followed by a high-voltage power Doppler image (20V). All Doppler images presented in the manuscript correspond to the 20V power Doppler acquisitions.

#### *In vivo* BURST imaging sequence with linear array

The same Verasonics Vantage system and L22-14vX transducer used for in vitro experiments was employed for in vivo imaging. The BURST pulse sequence began with a single low-pressure frame (transducer voltage: 1.6 V), followed by 16 high-pressure frames (transducer voltage: 20 V). To capture the entire depth of the brain, multiple transmit foci were used, with a BURST image acquired at each focus. These images were then summed to generate a composite BURST image. All images were captured in AM mode.

#### Doppler imaging with linear array

Power Doppler images of cerebral blood volume (CBV) were acquired at 15.625 MHz. The imaging protocol utilized a pulse sequence consisting of 15 tilted plane waves, varying from −14° to 14°, with a pulse repetition frequency of 500 Hz. A total of 200 coherently compounded frames were acquired. A singular value decomposition (SVD) clutter filter (cutoff = 40) was used to create the final power Doppler image showing CBV in the whole imaging plane^48^.

### In vivo image acquisition with matrix array

For volumetric BURST imaging, we used a modified version of Takoyaki BURST^21^ with a 15 MHz matrix array probe (Vermon), which transmits multiple focused beams and receives corresponding signals using a single group of 32 x 8 elements without parallel elements. Each volume image was acquired in AM mode. Therefore, 200 TX x 3 AM modes = 600 events were required to obtain one volume image. Transmit waveform parameters were the same as those used in in vitro experiments. The input voltage to the matrix array transducer was set to 1.6 V for low-pressure pulses and at least 25V for the high-pressure pulses.

Doppler imaging sequence with matrix array was based on previously established methods^21^.

### In vivo BURST image processing

Let *X*_1,_ *X*_2_, …, *X*_*N*_ be the sequence of *N* ultrasound images acquired following a pressure increase in the BURST scheme, with GV collapse occurring at the frame *X*_1_. To incorporate prior knowledge of the GV collapse signal’s temporal pattern, we computed the correlation between a template vector *Y* = [1 0 0 … 0] ∈ ℝ^*N*^ and the temporal profile of each voxel. The correlation coefficient *r* for voxel *i* ∈ {1, …, *k*}, where *k* is the number of voxels per frame, was calculated as,

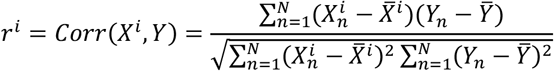

where 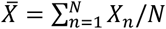 and 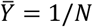 denote the temporal means of X and Y, respectively. To form the final BURST image, we retained only the voxels from the collapse frame *X*_1_ whose correlation coefficients are above a threshold *θ*:

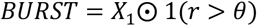

where ⨀ represents element-wise multiplication, and 1(·) is the indicator function. Although the choice of threshold *θ* is arbitrary, its value can be selected based on how aggressively we wish to exclude background voxels that lack GVs. Assuming there is no movement during image acquisition, voxels are independent, and each background voxel intensity follows an i.i.d. Gaussian process. The intensity of background voxel *i* can be expressed with random variables:

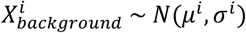

where *μ* is the mean intensity, and *σ* is the standard deviation of the intensity. Note that the superscript *i* implies that each background voxel may follow a different normal distribution. Given the independence of X and Y, and that at least one follows a univariate normal distribution, the sampling distribution of the correlation coefficient *r* is

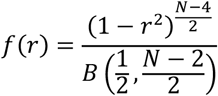

where B is the Beta function^49,50^. Since this distribution is not dependent of the voxel intensity parameters (*μ*^*i*^ and *σ*^*i*^), all background voxels share the same sampling distribution of *r*. Instead of using the above formula directly, we can take advantage of its relationship to the t-distribution^49,50^:

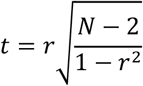

where *t* follows a t-distribution with *N* − *2* degrees of freedom. For 2D images, we chose *θ* = 0.7 (N = 16), corresponding to a background rejection rate of 0.9987.

During 3D ultrasound image acquisition, GVs collapse often spanned multiple frames. Suppose GVs collapse in frames *X*_1_, …, *X*_*a*_ and we are confident there is no further collapse from *X*_*b*_, where *a < b* ≤ *N*. In this case, we computed a BURST image from each of *X*_1_, …, *X*_*a*_ frames and sum them:

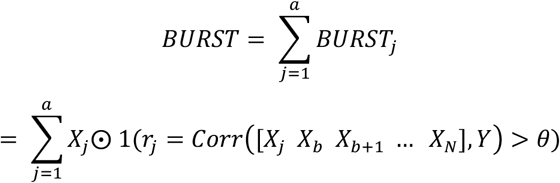

where *Y* = [1 0 0 … 0] ∈ ℝ^*N* −*b*+*2*^.. We chose *θ* = 0.7036 (N = 27, b = a+3 = 6), yielding a background rejection rate of 0.99991 for each *BURST*_*j*_.

## Data availability

All plasmids used in this study are available from M.G.S. under a material agreement with the California Institute of Technology. The key genetic constructs will be deposited with Addgene at the time of manuscript publication. Ultrasound acquisition and processing scripts used to generate key figures and results will be posted to a publicly accessible GitHub repository at the time of manuscript publication. Raw data are available from the corresponding author upon reasonable request.

## ACKNOWLEDGMENTS

The authors thank the laboratory of David Anderson for sharing their slide scanning microscope, Mickael Tanter for providing feedback on ultrasound post-processing methods, and Lindsey Salay, Joe Wekselblatt and Pierina Barturen for help with initial in vivo experiments. Confocal microscopy was performed in the Beckman Institute Biological Imaging Center with help from G. Spigolon. This research was supported by the National Institutes of Health (R01NS120828 and DP1EB033154 to M.G.S.). Related ongoing research is supported by the Advanced Research and Innovation Agency of the UK (9262914 to MGS). S.S. was supported by a National Science Foundation Graduate Research Fellowship Program fellowship. M.G.S. is an investigator of the Howard Hughes Medical Institute.

## CONTRIBUTIONS

S.S. conceived and planned the study, generated genetic constructs, and conducted in vitro and in vivo experiments. S.S. and K.Y.M.C. performed intracranial injections and cranial window surgeries. S.S., A.Y., and E.C.H conducted experiments in cultured neurons. I.U.H, and R.J.Z. conducted activity-dependent experiments in vitro. S.S. and S.L. conceptualized in vivo image post-processing method. S.S. conducted 3D imaging experiments. S.S. and J.R. performed histology and confocal imaging. C.R. helped establish in vivo ultrasound imaging scripts and post-processing methods. M.G.S. supervised the study. S.S. and M.G.S. wrote the manuscript. All other authors provided input on the final manuscript.

## COMPETING INTERESTS

S.S. and M.G.S. are inventors on one US patent application (US20230094152A1, published 30 March 2023) and additional unpublished application(s) related to this work filed by California Institute of Technology. The authors declare that they have no other competing financial interests.

## EXTENDED DATA FIGURES

**Extended Data Figure 1:**
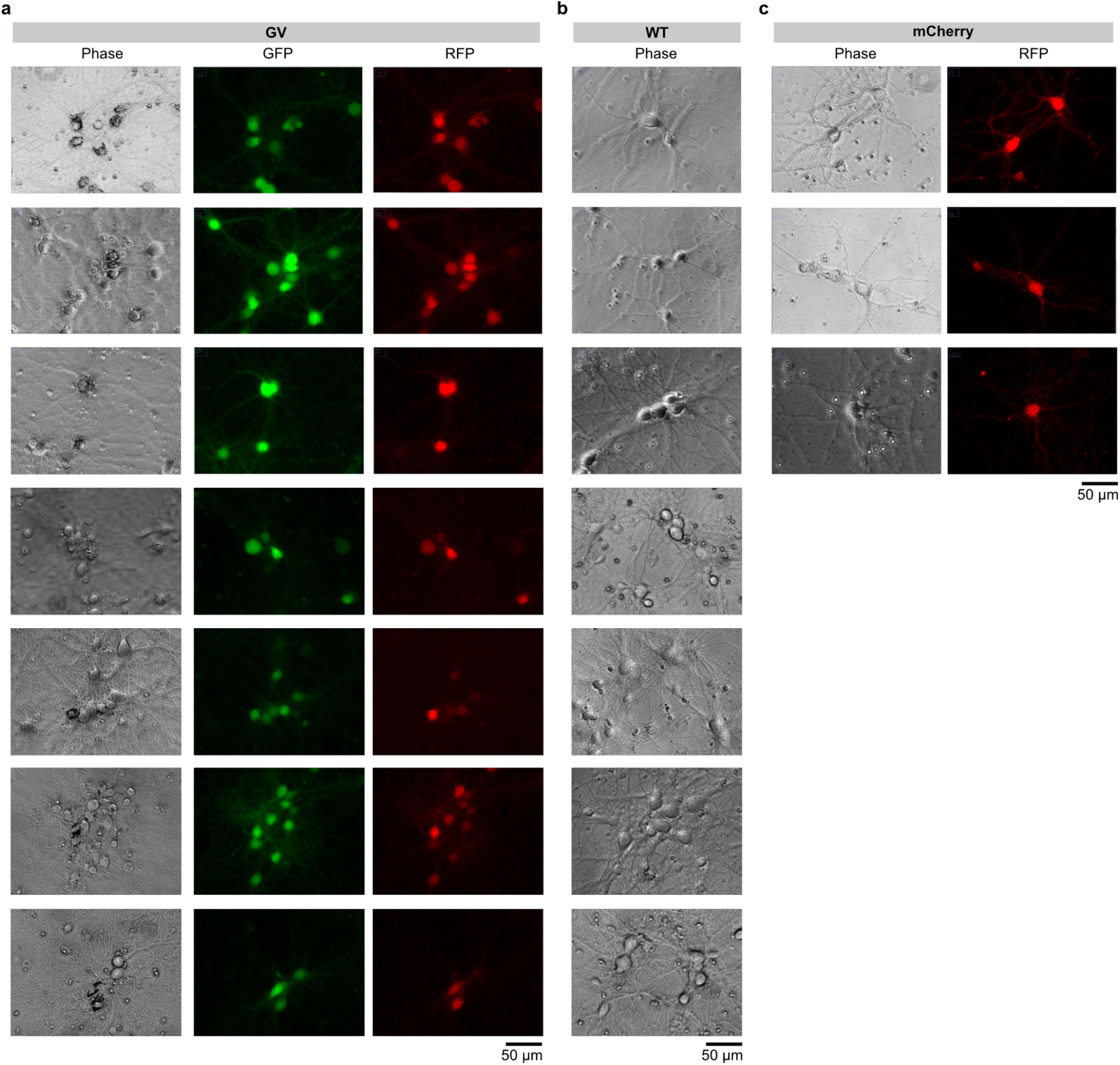
Optical microscopy of GV-expressing and control neurons. **a**, Representative phase contrast, GFP, and RFP fluorescence images of primary mouse neurons transduced with the GV-expressing AAV system. Clusters of GVs are present in cells with strong co-expression of GFP and RFP. **b**, Phase contrast images of WT untransduced neurons. **c**, Representative phase contrast and fluorescence images of neurons transduced with a constitutively expressing mCherry AAV. Cells show diffuse cytoplasmic expression of mCherry without clustered structures present in phase. Scale bars: 50 µm.

**Extended Data Figure 2:**
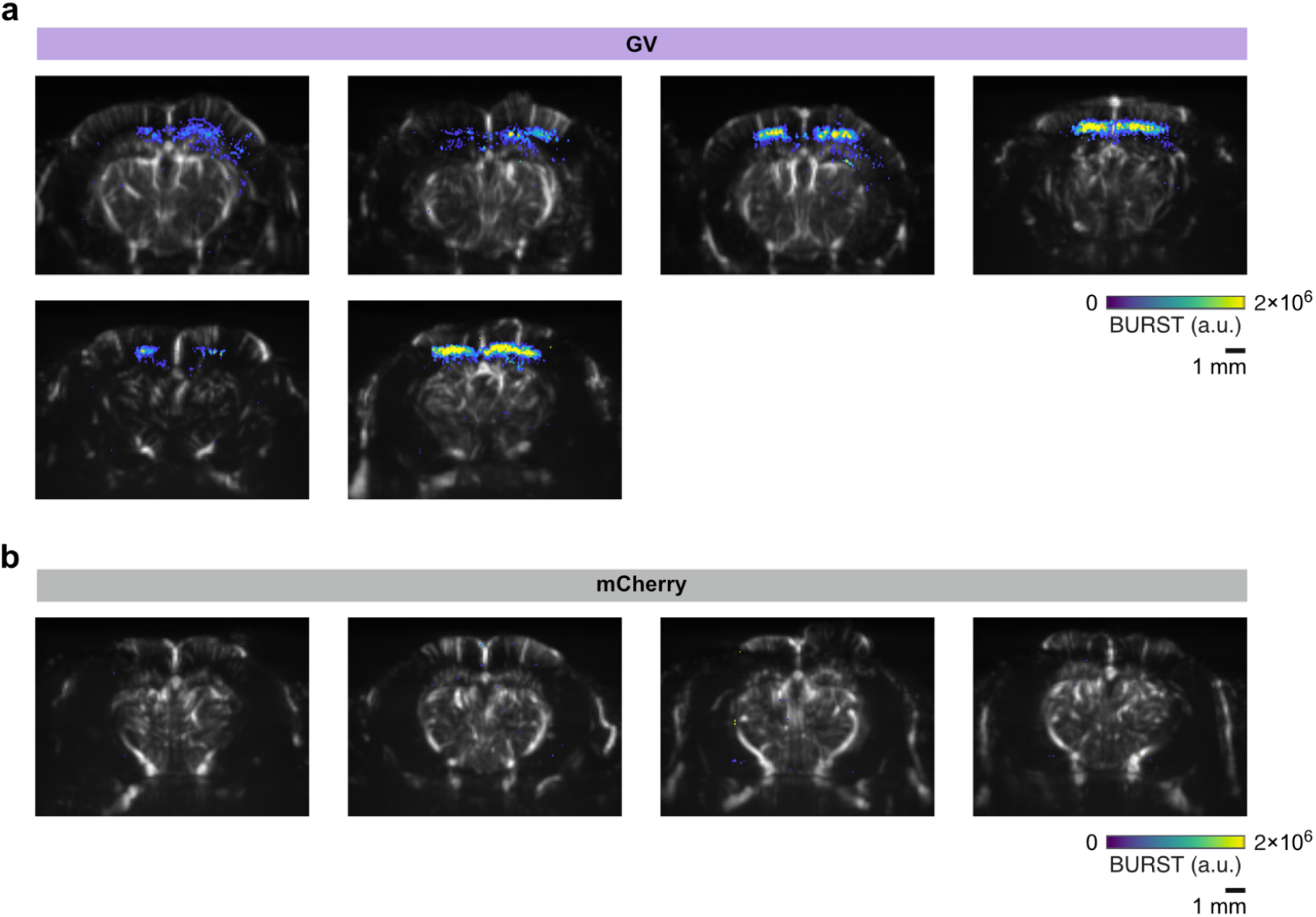
Ultrasound imaging of GV or mCherry expression in the mouse brain following intracranial AAV delivery. Representative BURST images (colorbar) of brains injected with **a**, the GV-expressing three vector AAVs or **b**, injected with a mCherry-expressing AAV, overlaid on top of Doppler images (grayscale). Each image is from a different mouse. The plane closest to the injection site, as determined by the Doppler image, and with the strongest BURST signal was selected from each mouse.

**Extended Data Figure 3:**
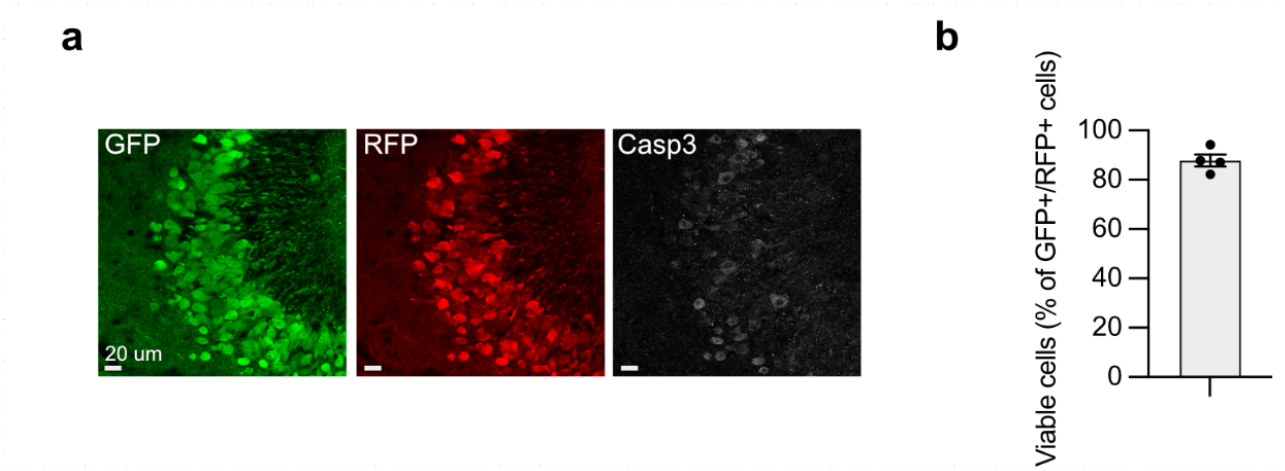
Assessment of cell viability in GV-expressing hippocampal neurons. **a**, Representative fluorescence images of the CA3 region from Dox-induced animals expressing GVs and imaged with the BURST sequence. Histological sections of the injection plane (AP –2.1 mm), showing GFP and RFP expression alongside cleaved caspase-3 (Casp3) staining. CA3 was the hippocampal subregion with the highest observed level of cell death. **b**, Quantification of Casp3^+^ cells as a percentage of GFP^+^ and RFP^+^ cells. n = 4 mice. n = 111 ± 34 (mean ± s.d.) triple-transduced cells per mouse. Scale bar: 20 μm.

**Extended Data Figure 4:**
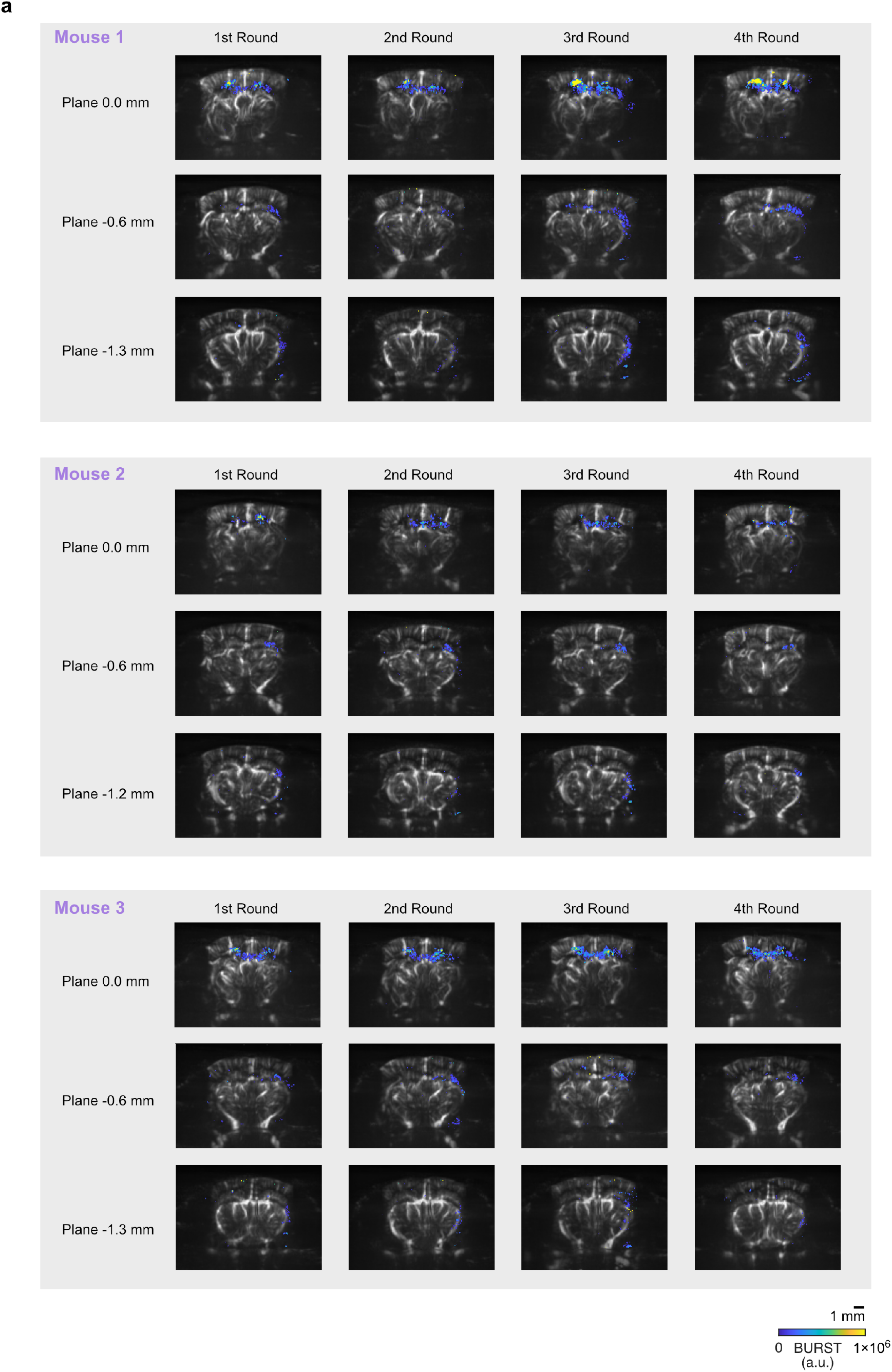
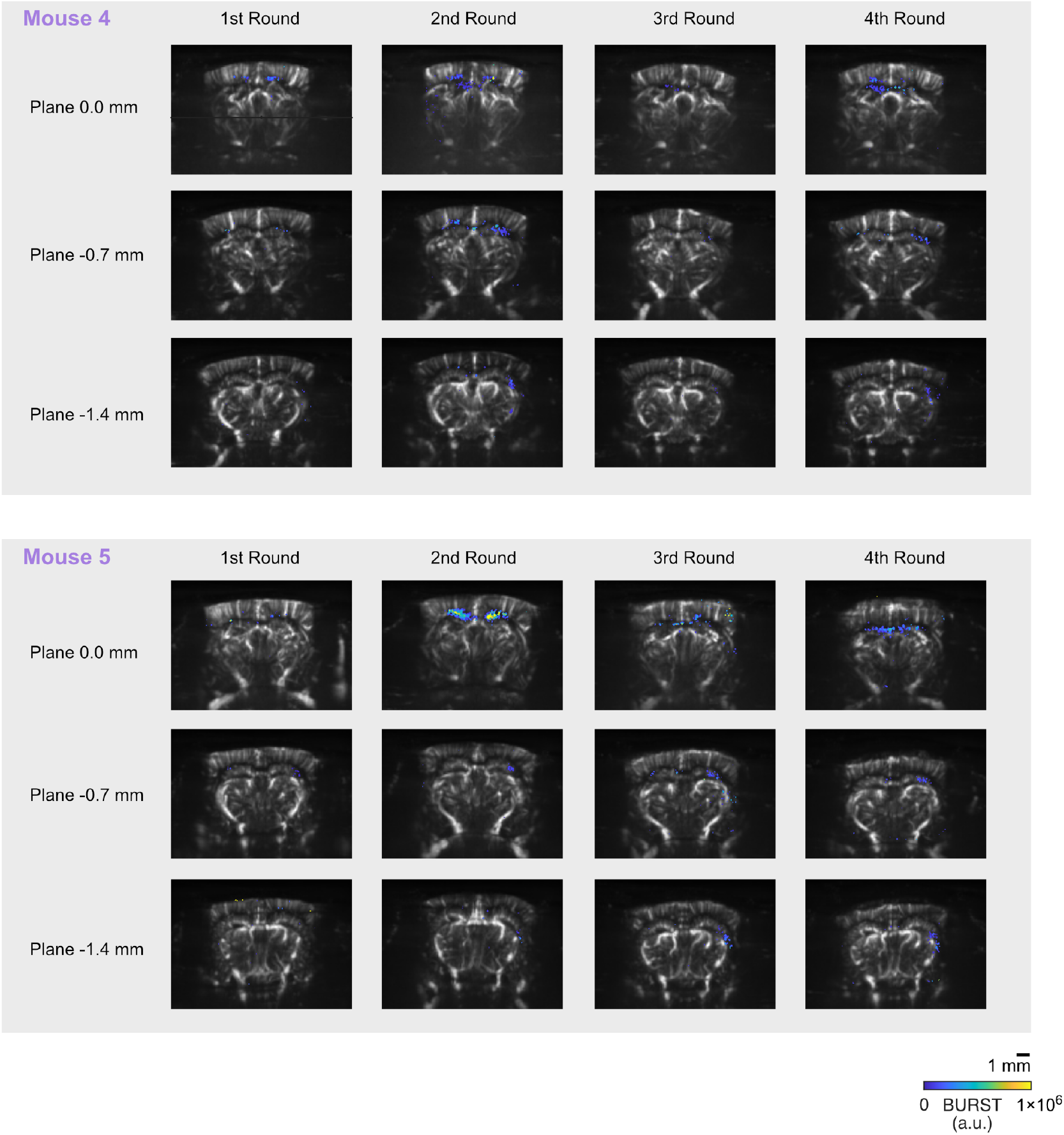
Ultrasound imaging of repeated GV expression in the brain. **a**, Ultrasound images of repeated GV expression in mice implanted with an acoustically transparent cranial window. Mice received Dox for 48 hours prior to each imaging session, with induction-imaging cycles spaced two to three weeks apart. For quantification, the imaging plane with the strongest BURST signal from the first session was used for all timepoints. BURST images are shown in color and overlaid on grayscale Doppler anatomical images. n = 5 mice.

**Extended Data Figure 5:**
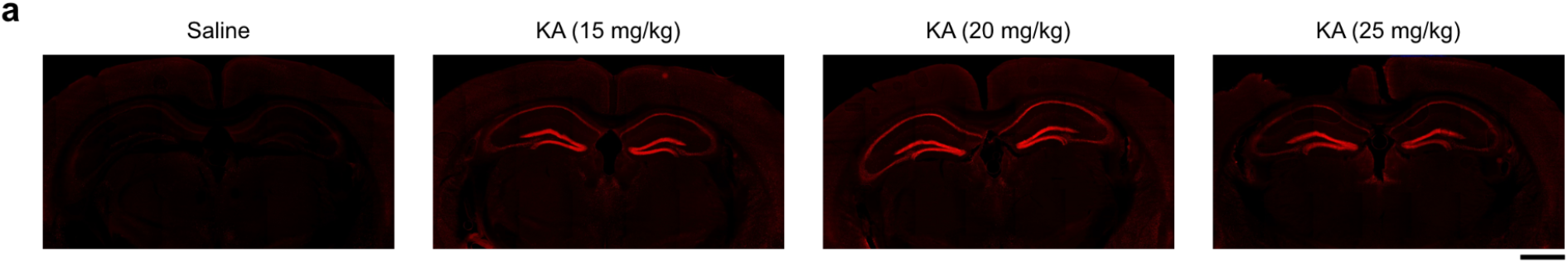
Systemic kainic acid injection induces cFos expression in the hippocampus. **a**, Representative images of cFos immunostaining in coronal brain sections from mice injected intraperitoneally with either saline or varying doses of kainic acid. Mice were sacrificed 30 minutes after injection. Scale bar: 1 mm.

**Extended Data Figure 6:**
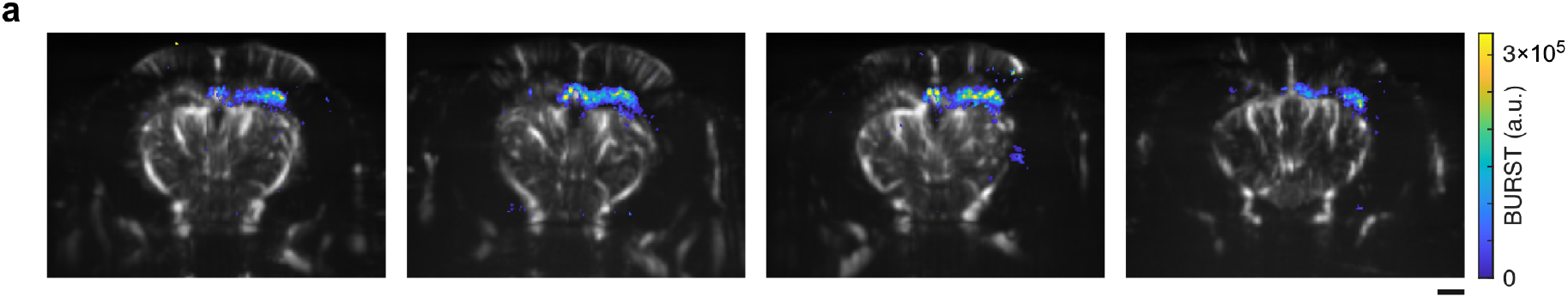
Co-expression of a constitutive fluorophore with Tet-inducible *gvpA* does not affect GV expression in the hippocampus. **a**, Representative ultrasound images of the hippocampus from mice injected with an AAV encoding a Tet-inducible *gvpA* and a constitutive iRFP670 fluorophore (as in **Fig. 5a**), along with the two AAVs encoding assembly factors (as in **Fig. 2a**). BURST signal is shown in color and overlaid on grayscale Doppler images. n = 4 mice. Scale bar: 1 mm.

**Extended Data Figure 7:**
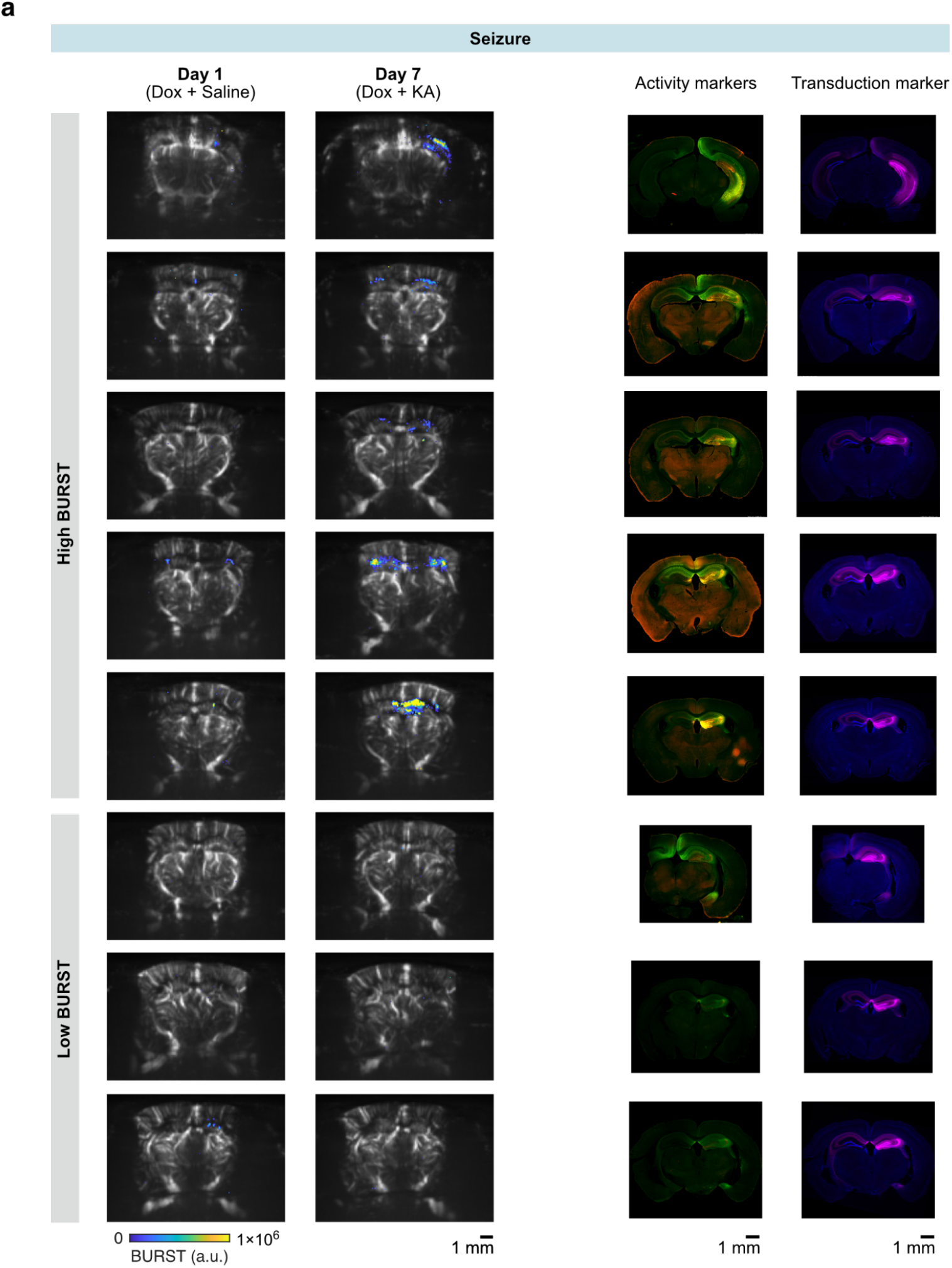
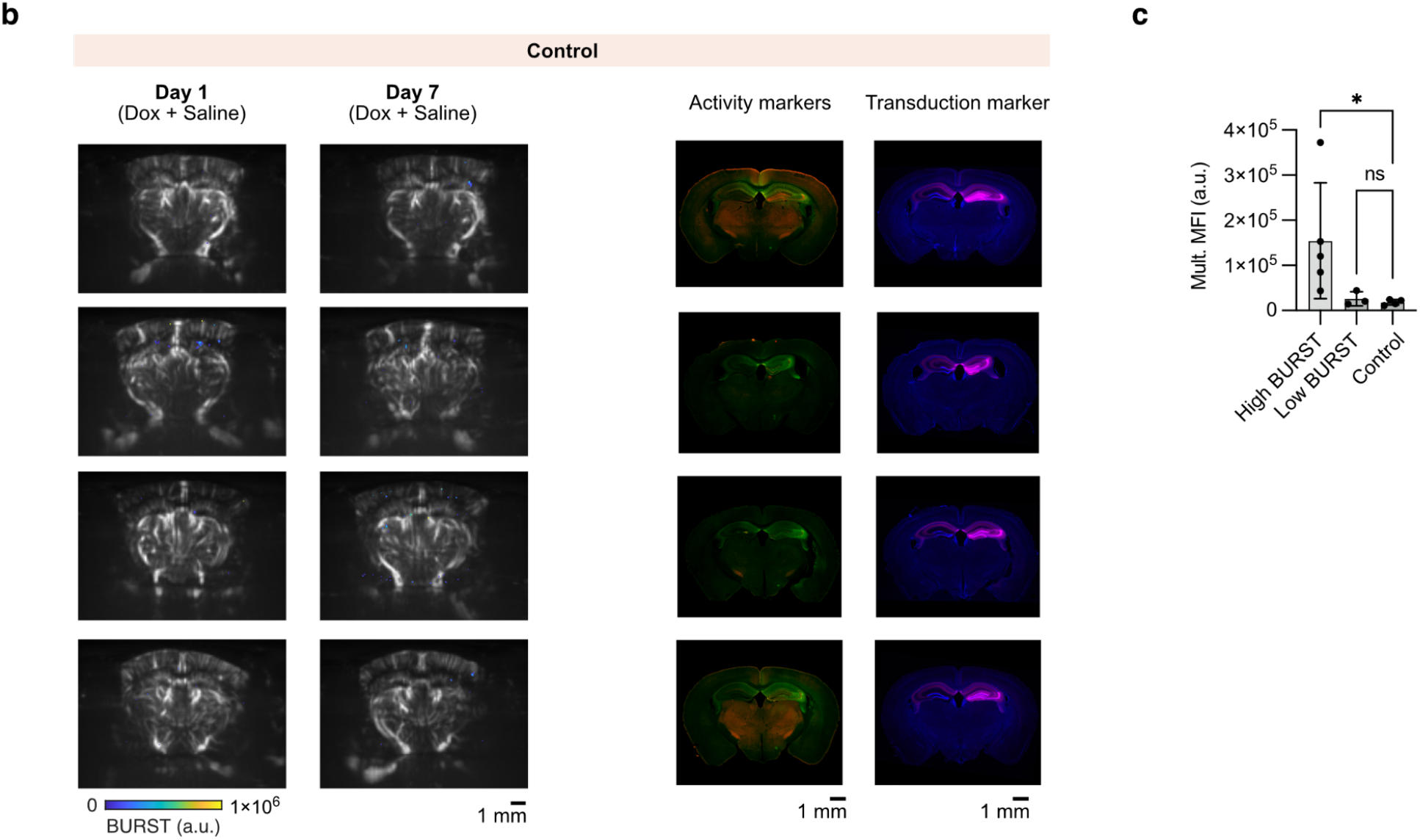
Ultrasound imaging and fluorescence microscopy of reporter expression in mice with and without induced seizures. **a**, Representative ultrasound and histological images from seizure-induced and **b**, control mice. Left panels: BURST images (colormap) overlaid on grayscale Doppler anatomical images acquired on day 1 (baseline, post-Dox/saline) and day 7 (post-Dox/KA or Dox/saline). Right panels: GFP and RFP fluorescence showing IEG-driven reporter expression (activity markers), and iRFP670 (transduction marker) with DAPI staining. Several seizure-induced mice did not show a significant increase in BURST signal on day 7 (Low BURST), consistent with weak fluorescence expression. For seizure-induced mice, representative imaging planes were selected based on proximity to the injection site and the strongest observed change in BURST signal between day 1 and day 7. **c**, Product of GFP and RFP channel mean fluorescence intensities in the right hippocampus normalized to the contralateral side (Mult. MFI). High BURST: Seizure-induced mice exhibiting high BURST signal, n = 5. Low BURST signal: Seizure-induced mice exhibiting low BURST signal, n = 3. Control, n = 4. One-way ANOVA without correction for multiple comparisons. P-values from left to right: 0.0417, 0.904. n.s. p ≥ 0.5, *p < 0.05, **p < 0.01, ***p < 0.001. Scale bars: 1 mm. KA: Kainic Acid.

**Supplementary Table 1.**
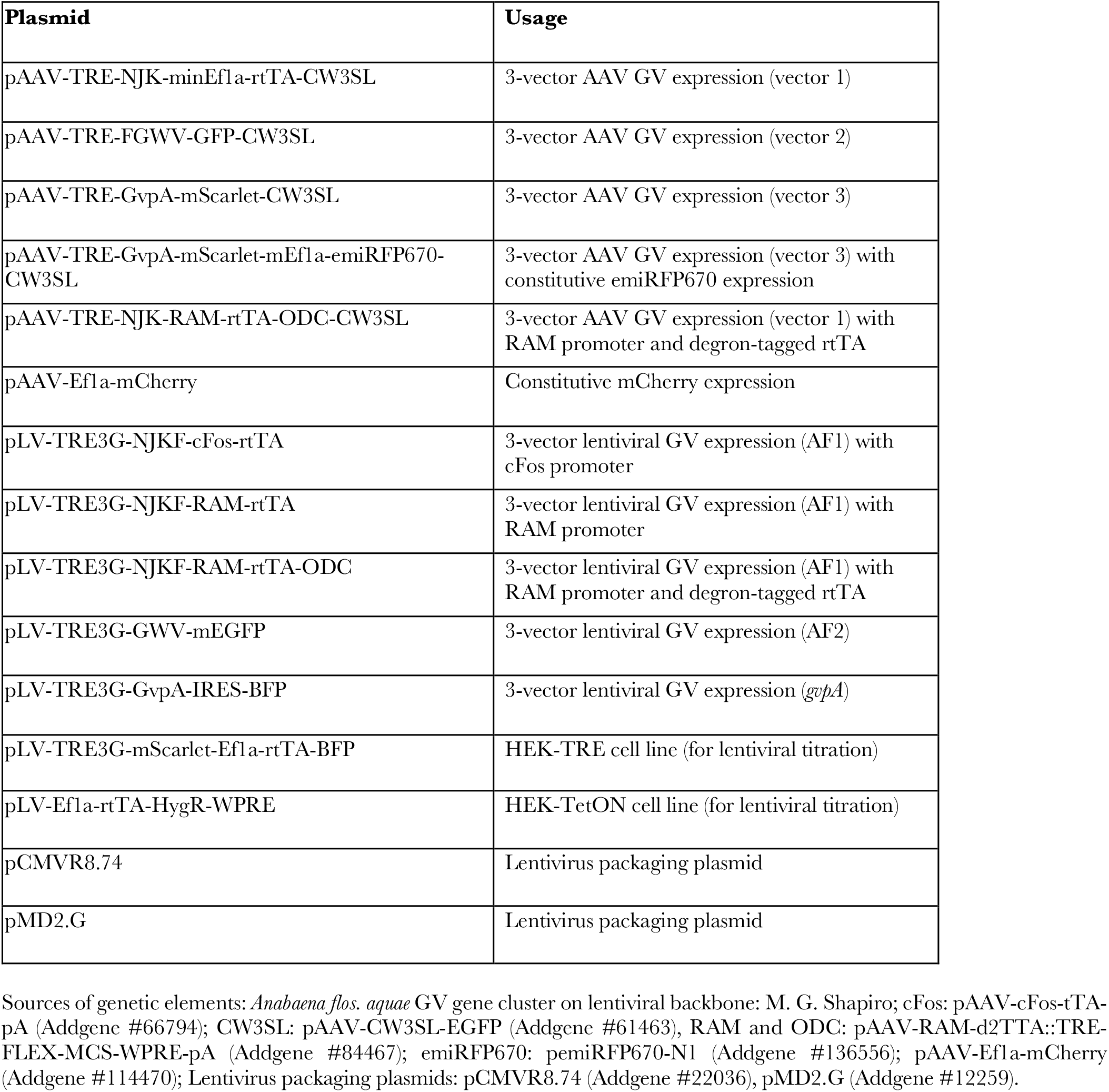
Genetic constructs used in the study. Sources of genetic elements: *Anabaena flos. aquae* GV gene cluster on lentiviral backbone: M. G. Shapiro; cFos: pAAV-cFos-tTA-pA (Addgene #66794); CW3SL: pAAV-CW3SL-EGFP (Addgene #61463), RAM and ODC: pAAV-RAM-d2TTA::TRE-FLEX-MCS-WPRE-pA (Addgene #84467); emiRFP670: pemiRFP670-N1 (Addgene #136556); pAAV-Ef1a-mCherry (Addgene #114470); Lentivirus packaging plasmids: pCMVR8.74 (Addgene #22036), pMD2.G (Addgene #12259).

